# Human anterolateral entorhinal cortex volumes are associated with preclinical cognitive decline

**DOI:** 10.1101/080374

**Authors:** RK Olsen, L-K Yeung, A Noly-Gandon, MC D’Angelo, A Kacollja, VM Smith, JD Ryan, MD Barense

**Author notes:** Current address: Taub Institute Columbia University Medical Center 630 West 168th Street P&S Box 16 New York, NY 10032. Corresponding authors’ information: Rosanna K. Olsen 3560 Bathurst Street Toronto, ON M6A 2E1 Lok-Kin Yeung 630 West 168th Street New York, NY 10032.

## Abstract

We investigated whether older adults without subjective memory complaints, but who present with cognitive decline in the laboratory, demonstrate atrophy in medial temporal lobe (MTL) subregions associated with Alzheimer's disease. Forty community-dwelling older adults were categorized based on Montreal Cognitive Assessment (MoCA) performance. Total grey/white matter, cerebrospinal fluid, and white matter hyperintensity load were quantified from whole-brain T1-weighted and FLAIR magnetic resonance imaging scans, while hippocampal subfields and MTL cortical subregion volumes (CA1, dentate gyrus/CA2/3, subiculum, anterolateral and posteromedial entorhinal, perirhinal, and parahippocampal cortices) were quantified using high-resolution T2-weighted scans. Cognitive status was evaluated using standard neuropsychological assessments. No significant differences were found in the whole-brain measures. However, MTL volumetry revealed that anterolateral entorhinal cortex (alERC) volume -- the same region in which Alzheimer's pathology originates -- was strongly associated with MoCA performance. This is the first study to demonstrate that alERC volume is related to cognitive decline in preclinical, community-dwelling older adults.

## 1 Introduction

Alzheimer’s disease (AD) is a devastating neurodegenerative illness with widespread societal and economic consequences. Due to the progressive nature of the disease, early and effective diagnosis of AD is critical for the development and delivery of drug treatments and/or interventions. Pathological changes in the medial temporal lobe (MTL) may occur several years prior to the onset of subjective memory complaints and diagnosis in the clinic (Sperling et al., 2011). The goal of the current work is to identify structural MTL measurements that indicate AD susceptibility in an ostensibly cognitively healthy community sample of older adults. Critically, in addition to the hippocampal subfields, this study investigated MTL cortex subregions that develop AD pathology (cellular loss, tau abnormalities, and tangle pathology) at the earliest stage of the disease (Jack and Holtzman, 2013; Jack et al., 2013, 2010).

Neuropsychological testing is used for the diagnosis of Mild Cognitive Impairment (MCI), a condition which often progresses to AD (Petersen, 2004; Sperling et al., 2011). The Montreal Cognitive Assessment (MoCA) is a brief screening tool that demonstrates excellent sensitivity and specificity in detecting MCI (Markwick et al., 2012; Nasreddine et al., 2005) and predicting conversion to AD (Julayanont et al., 2014). Older adults who performed poorly on the MoCA also exhibited cognitive impairments in memory (D’Angelo et al., 2016; Yeung et al., 2013), have shown abnormal electrophysiological signatures (Newsome et al., 2013), and demonstrated similarly low performance on visual discrimination tasks as MCI patients (Newsome et al., 2012). The current investigation used detailed volumetric analyses to investigate brain atrophy associated with poor MoCA performance in a group of community-dwelling older adults who, critically, have no current subjective memory complaints and no MCI diagnosis.

Advanced AD is associated with distributed neocortical structural changes (Scahill et al., 2002; Thompson et al., 2003); however the earliest stages of the disease are thought to develop within the MTL (Braak and Braak, 1991). Specific subregions of the MTL, including the entorhinal cortex (ERC; Gómez-Isla et al., 1996; Krumm et al., 2016; Whitwell et al., 2007) and the CA1 subfield of the hippocampus (Chételat et al., 2008; de Flores et al., 2015; Gerardin et al., 2009; Iglesias et al., 2015; Kerchner et al., 2012; La Joie et al., 2013; Mueller and Weiner, 2009; Mueller et al., 2010a; Pluta et al., 2012; Tang et al., 2014; Yassa et al., 2010; Yushkevich et al., 2015b) exhibit volumetric decreases in individuals with MCI.

The anterolateral portion of entorhinal cortex (alERC) and the perirhinal cortex (PRC) were recently identified as a primary sites of cerebral blood volume (CBV) reductions in a group of 12 humans who subsequently developed AD (Khan et al., 2014). Similar CBV reductions were observed in the corresponding regions in transgenic mouse models of AD, suggesting that the alERC and the PRC are affected earliest by AD pathology (Khan et al., 2014). Asymptomatic individuals with AD genetic risk were shown to have reductions in hippocampal and MTL cortical subregions, including the ERC, indicating that disease-related structural atrophy can precede subjective memory complaints (Fox et al., 1996; Harrison et al., 2016). However, to our knowledge, there has been no investigation of volumetric changes to the alERC in asymptomatic individuals; here we provide the first study to employ manual segmentation of the alERC as a distinct region from the ERC as a whole.

We hypothesized that older adults who reported no subjective memory impairments, but nonetheless scored below the recommended cutoff MoCA score (<26) would have reduced volume in the MTL, specifically within the alERC, PRC, and the CA1 subfield of the hippocampus. To our knowledge only two studies have examined the relationship between MoCA performance and MTL volumes, and these studies focused on the hippocampus proper (Gupta et al., 2015; Paul et al., 2011), which develops AD pathology later than the adjacent alERC (Braak and Braak, 1991). This is the first study to address the relationship between alERC volume and cognitive status. While the primary investigation focused on the MTL, global estimates of brain volume (total grey matter, white matter, and cerebrospinal fluid) were also investigated (Gupta et al., 2015). Finally, to rule out undetected stroke and investigate potential contributions of vascular pathology to cognitive impairment, volumetric assessment of white matter hyperintensities (WMH) was conducted (Brickman et al., 2015).

## 2 Material and methods

### 2.1 Participants

Forty community-dwelling, older adult participants (30 female; mean age=71.4 years, range=59-81, mean education=16.3 years, range=12-23) were recruited from participant databases at the RRI/Baycrest and the University of Toronto. All participants received the MoCA (Nasreddine et al., 2005), and were selected to create two age-matched groups that differed solely on the basis of their MoCA score. A score of 26 is the recommended threshold score for primary care physicians to provide further dementia screening (Damian et al., 2011); thus, the two groups were defined as an “at-risk” group (17 female; mean age=72.5 years, range=59-81, mean education=16.2 years, range=12-22) who scored below 26 (indicating potentially pathological cognitive impairment and risk for MCI, mean score=23.4, range=17-25), and a “healthy” group (13 female; mean age=70.3 years, range=63-77, mean education=16.6 years, range=12-23) who scored 26 and above (mean score=27.9, range=26-30). T-tests showed no difference between the two groups in age, t(38)=1.29, p=.20, or years of education, t(38)=.51, p=.61, but a significant difference in MoCA score, t(38)=7.87, p<.001.

All participants were fluent English speakers with normal or corrected-to-normal vision, and were screened for non-MRI compatible metal implants, color blindness, diabetes, neurological disorders, stroke, or brain trauma. All participants were informed about the nature of the experiment and its risks, and gave written informed consent. The Research Ethics Board of the University of Toronto and the RRI approved this research. All participants received monetary compensation for participation, following standard practices at the RRI.

### 2.2 Neuropsychological Battery

All participants received a battery of neuropsychological tests (Osterrieth, 1944; Reitan and Wolfson, 1985; Warrington and James, 1991; Wechsler, 2009, 1999; Wechsler et al., 2008) to characterize cognitive performance (Table 1). The magnitude of subjective memory complaints in everyday memory functioning was also quantified to evaluate whether participants in either group had self-awareness of memory difficulties (Gilewski et al., 1990).

**Table 1.** Neuropsychological Battery.

| Test | Healthy<br>Older Adults | At-risk<br>Older Adults | Effect size of<br>difference<br>between groups<br>(Cohen's d) |
| --- | --- | --- | --- |
| MoCA (/30)** | 27.9 (1.7)<br>Normal range | 23.4 (1.9)<br>Impaired | 2.49 |
| Visuospatial/Executive (/5)<sup>†</sup> | 4.2 (1.0) | 3.7 (0.9) | 0.60 |
| Naming (/3)** | 3.0 (0.2) | 2.4 (0.7) | 1.09 |
| Attention (/6)* | 5.9 (0.4) | 5.3 (0.9) | 0.78 |
| Language (/3) | 2.8 (0.4) | 2.4 (0.8) | 0.57 |
| Abstraction (/2) | 1.9 (0.3) | 1.8 (0.6) | 0.33 |
| Memory (/5)** | 4.1 (1.1) | 2.1 (1.4) | 1.57 |
| Orientation (/6)* | 6.0 (0.0) | 5.8 (0.6) | 0.64 |
| WMS-IV LM Immediate Recall<br>Scaled Score (/20) | 11.9 (2.9)<br>70.6%ile | 10.9 (2.3)<br>58.7%ile | 0.40 |
| WMS-IV LM Delayed Recall<br>Scaled Score (/20)* | 11.8 (2.5)<br>68.3%ile | 9.9 (2.1)<br>49.7%ile | 0.83 |
| WMS-IV LM Recognition<br>Accuracy** | 86% (10%) | 78% (9%) | 0.88 |
| Trails A | 42.7s (11.6s)<br>39.2%ile | 43.6s (15.5s)<br>43.1%ile | 0.07 |
| Trails B* | 79.0s (30.5s)<br>63.3%ile | 102.1s (36.7s)<br>51.7%ile | 0.69 |
| Digit Span Forward Score<sup>†</sup> (/16) | 10.9 (2.0)<br>61.1%ile | 9.6 (2.2)<br>44.1%ile | 0.60 |
| Digit Span Backward Score*<br>(/14) | 7.6 (2.2)<br>41.5%ile | 6.0 (2.5)<br>24.5%ile | 0.65 |
| Rey-Osterrieth Complex Figure |  |  |  |
| Copy (/32) | 26.8 (5.2)<br>26.3%ile | 27.3 (5.9)<br>30.7%ile | 0.09 |
| Immediate Recall (/32) | 13.0 (6.8)<br>43.7%ile | 11.9 (6.6)<br>39.4%ile | 0.18 |
| Delayed Recall (/32) | 12.0 (6.6)<br>39.3%ile | 9.9 (6.4)<br>31.1%ile | 0.32 |
| WASI Vocabulary (/80)<sup>†</sup> | 62.3 (5.3)<br>74.9%ile | 56.9 (12.1)<br>58.4%ile | 0.58 |
| WASI Similarities (/48)<sup>†</sup> | 37.7 (3.8)<br>80.2%ile | 34.9 (5.6)<br>70.4%ile | 0.60 |
| WASI Matrix Reasoning (/32)** | 24.8 (4.7)<br>84%ile | 19.4(7.5)<br>68.1%ile | 0.86 |
| WASI Block Design (/71) | 33.3 (15.1)<br>57.3%ile | 27.8 (14.3)<br>52.7%ile | 0.38 |
| VOSP Shape Detection (/20)<br>(Cut-off score < 15) | 19.4 (0.9)<br>Pass | 19.0 (1.3)<br>Pass | 0.31 |
| VOSP Incomplete Letters (/20)*<br>(Cut-off score < 16) | 19.6 (0.8)<br>Pass | 19.0 (0.8)<br>Pass | 0.70 |
| VOSP Dot Counting (/10) (Cut-off score < 8) | 9.9 (0/3)<br>Pass | 9.7 (0.5)<br>Pass | 0.42 |
| VOSP Position Discrimination (/20)<sup>†</sup> (Cut-off score < 18) | 19.7 (0.6)<br>Pass | 18.9 (2.1)<br>Pass | 0.54 |
| VOSP Number Location (/10)*<br>(Cut-off score < 7) | 9.7 (0.7)<br>Pass | 8.6 (2.0)<br>Pass | 0.70 |
| VOSP Cube Analysis (/10) (Cut-off score < 6) | 9.7 (0.7)<br>Pass | 9.2 (1.6)<br>Pass | 0.40 |
| VOSP Silhouettes (/30) (Cut-off score < 15) | 20.2 (5.2)<br>Pass | 19.8 (5.3)<br>Pass | 0.08 |
| VOSP Object Decision (/20)<br>(Cut-off score < 14) | 17.2 (1.9)<br>Pass | 16.4 (2.0)<br>Pass | 0.40 |
| VOSP Progressive Silhouettes (/20) (Cut-off score > 15) | 10.0 (2.6)<br>Pass | 10.4 (3.5)<br>Pass | 0.12 |
| Subjective memory rating<br>(Memory functioning questionnaire, /448) | 295.9 (68.9)<br>Minimal subjective difficulties | 289.7 (38.7)<br>Minimal subjective difficulties | 0.11 |

**Note.** Mean and standard deviation are listed for each group. Maximum and cut-off scores for tests indicated in parentheses in left column. WMS-IV LM = Wechsler Memory Scale, 4<sup>th</sup> Edition, Logical Memory subtest; WASI = Weschler Abbreviated Scale of Intelligence; VOSP = Visual Object and Spatial Perception battery. <sup>†</sup> indicates a trend towards significant difference between healthy and at-risk older adults at $p < .10$ , \* indicates a significant difference at $p < .05$ , \*\* $p < .01$ . All t-tests were two-tailed.

Neuropsychological testing was completed in a separate session prior to the MRI scan (*M*=3 months; *SD*=5.5 months). Results are presented in Table 1 along with effect sizes for the group differences. Both groups of participants performed in the low-average to high-average range on neuropsychological tests of delayed memory, working memory, executive function, semantic memory, and visuospatial perception. However, when directly comparing the at-risk group to the healthy group, medium to large effects sizes on several standardized tests of delayed memory were observed. These included the WMS Logical Memory tests (delayed recall and recognition) and the Rey–Osterrieth delayed recall test. The at-risk group also demonstrated lower scores on tests of working memory (WAIS digit span), executive function (Trails A & B), and semantic memory (WASI vocabulary). Visuospatial performance was largely intact. Scores on the subjective memory functioning questionnaire were equivalent among the groups, and the scores obtained indicated that neither group self-reported significant functional memory difficulties outside of the laboratory. Although the at-risk individuals performed below the MoCA cut-off, they did not present with subjective memory complaints, and performed in the average range on these standard neuropsychological tests; therefore, they do not meet the criteria for MCI. Petersen, 2004).

### 2.3 Structural Image Acquisition

All neuroimaging was done on a 3T Siemens Trio scanner using a 12-channel head coil.

Participants received a T1-weighted magnetization-prepared, rapid acquisition with gradient echo image (MP-RAGE) whole-brain anatomical scan (TE/TR=2.63 ms/2000ms, 160 axial slices perpendicular to the AC-PC line, 256x192 acquisition matrix, voxel size=1x1x1 mm, FOV=256mm). The MP-RAGE scan was used to obtain measures of brain and head size, as well as for the quantification of global grey and white matter and cerebral spinal fluid (CSF). The MP-RAGE scan was also used for slice placement during the acquisition of a subsequent high-resolution T2-weighted scan in an oblique-coronal plane, perpendicular to the hippocampal long axis (TE/TR=68ms/3000ms, 20-28 slices depending on head size, 512x512 acquisition matrix, voxel size=0.43x0.43x3 mm, no skip, FOV=220mm). For the high-resolution, T2-weighted scan, the first slice was placed anterior of the collateral sulcus (including the temporal pole where possible), and the last slice placed posterior to the hippocampal tail, to ensure coverage of the entire hippocampus and MTL cortices. A whole-brain fluid-attenuated inversion recovery image (FLAIR; TE/TR=97/9000ms, 30-32 axial slices perpendicular to the AC-PC line, voxel size=0.875x0.875x5mm, 212x256 acquisition matrix, FOV=220mm, TI=2500ms) was collected to check for the presence of strokes and WMH.

### 2.4 Global Brain Measure Estimation Using Automated Segmentation

Global estimates of cortical grey and white matter volume, CSF, and the estimated total intracranial Volume (eTIV) were obtained using FreeSurfer (version 5.3; http://surfer.nmr.mgh.harvard.edu/). The eTIV was used to correct MTL subregion and WMH volumes for head-size (as a proxy for intracranial volume;1 Buckner et al., 2004). The technical details of the volumetric segmentation procedures are described by Fischl and colleagues (Fischl et al., 2002).

WMH load was estimated using the LST toolbox version 1.2.3 (http://www.applied-statistics.de/lst.html), an automated tool for the segmentation of T2-hyperintense WMH in FLAIR images (Schmidt et al., 2012), which has recently been used to evaluate WMH load in patients diagnosed with probable AD (Morgen et al., 2015). We employed the lesion growth algorithm, which operates in native space using the following steps. First, FLAIR images were bias-corrected to remove MRI field inhomogeneities, next FLAIR images were coregistered to T1-weighted images and each tissue class (grey matter, white matter, CSF) was determined from the T1-weighted images. The distribution of FLAIR intensities for each tissue was then analyzed with the aim of detecting hyperintense outliers, indicating lesion voxels. According to their spatial location, the lesion voxels were categorized in three lesions belief maps (grey and white matter, CSF), which were summed into a single lesion belief map. This initial (conservative) lesion map, was set as a binary version of the GM belief map on which the default kappa threshold was applied (k=.3). Visual inspection of the lesion probability maps and their corresponding FLAIR images confirmed that the default kappa threshold was optimal for the current data. Finally, the lesion growth algorithm refined the lesion probability map as neighbouring voxels were iteratively analyzed and assigned to white matter, grey matter or lesion until no further voxels are assigned to lesions. The result was a lesion probability map for each subject that was transformed into binary maps using a threshold of .5. WMH volumes were then extracted from the binary maps.

### 2.5 Manual Segmentation of the MTL Subregions

Manual segmentation was performed on the T2-weighted images, in participants’ native space, on the oblique-coronal plane perpendicular to the long axis of the hippocampus (Figure 1; in-plane resolution=0.43x0.43mm). A single rater (L.Y.), who was blind to MoCA score/group status, manually delineated three hippocampal subfields (CA1, a region combining dentate gyrus, CA2 and CA3 [DG/CA2/3], subiculum), and four MTL cortex subregions (alERC, posteromedial entorhinal (pmERC), PRC, and parahippocampal cortices (PHC)) in FSLview. A second rater (R.K.O.), who was also blind to MoCA score/group status, segmented the same regions to provide an index of inter-rater reliability (see below). This segmentation protocol is largely similar to the Olsen-Amaral-Palombo (OAP) protocol which has been used for previous volumetric investigations of the MTL (Olsen et al., 2013, 2009; Palombo et al., 2013; Yushkevich et al., 2015a). The OAP protocol typically includes two additional regions of interest, which cover the anterior head and the posterior tail of the hippocampus. However, in the current older adult cohort, the outer definition of the anterior and posterior slices was difficult and unreliable; thus, these regions were excluded from further analysis. The segmentation of the hippocampal subfields followed published anatomical atlases (Amaral and Insausti, 1990; Duvernoy, 2005). The segmentation of the MTL cortices followed the protocol of Insausti and colleauges for the ERC and PRC, and the protocol of Pruessner and colleagues for the PHC (Insausti et al., 1998; Pruessner et al., 2002). The subdivision of the entorhinal cortex into alERC and pmERC following the protocol of Maas, Berron and colleagues (Maass et al., 2015).

**Figure 1.**
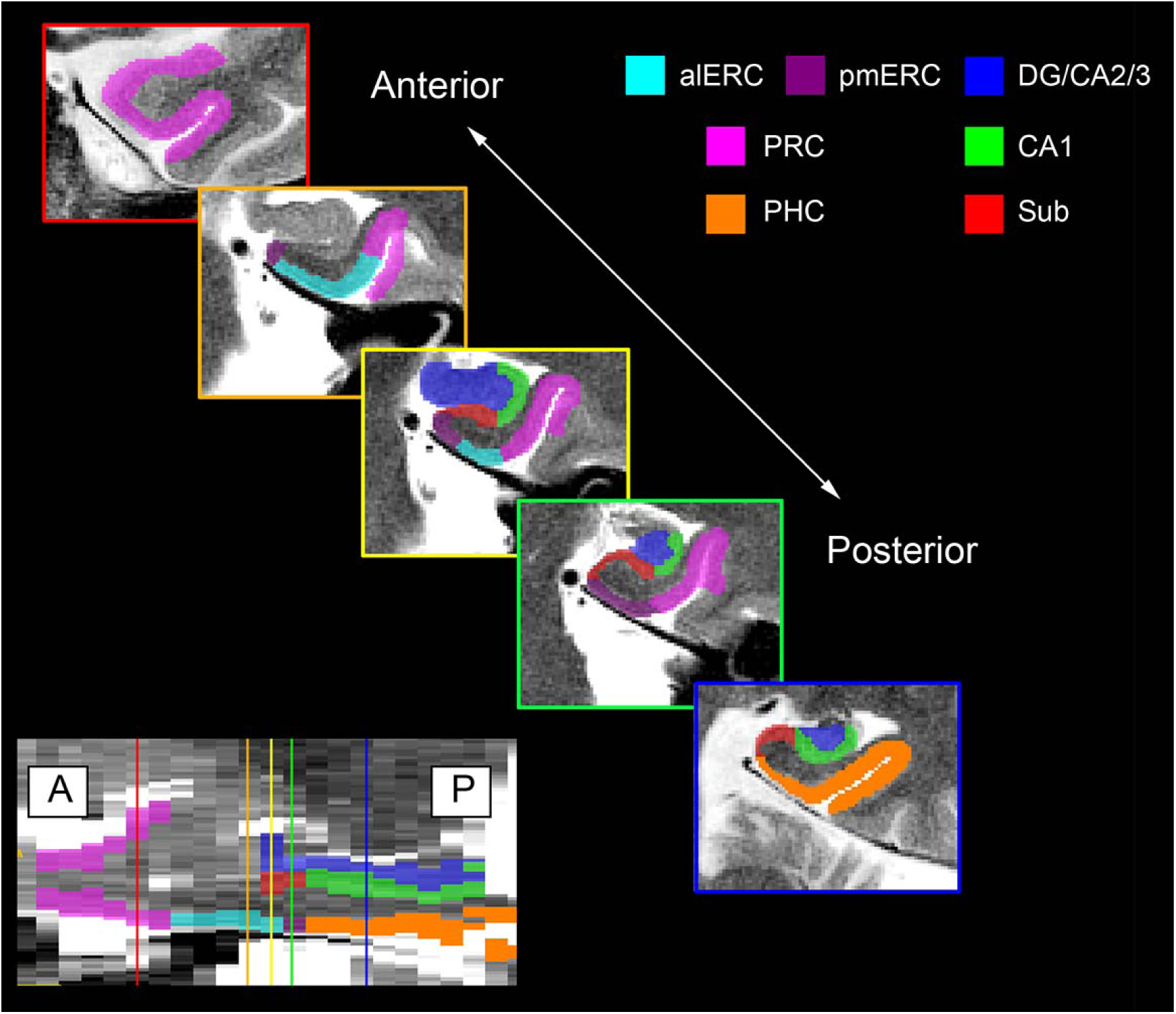
Segmentation protocol. Example coronal slices, spanning the anterior-posterior axis of the MTL. Slices depict the manual segmentation protocol used in the current study and regions of interest depict each of the seven regions (3 hippocampal subfields, 4 MTL cortex subregions) that were compared across groups. Note that the segmentation of the hippocampal subfields, perirhinal and parahippocampal cortices is the same protocol used in previous work (Olsen et al., 2013). The segmentation of the entorhinal cortex into anterolateral and posteromedial segments follows the protocol of Maass, Berron and colleagues (2015).

### 2.6 MTL Manual Segmentation Reliability

Intra-rater reliability was established by comparing segmentation of five randomly selected scans, completed by the same rater (L.Y.) after a delay of 1-4 months. Inter-rater reliability was evaluated by comparing the segmentation of five randomly selected scans by a second rater (R.K.O) to those of L.Y. Reliability was assessed using the Intra-class correlation coefficient (ICC, which evaluates volume reliability) and the Dice metric (which also takes spatial overlap into account), computed separately for each region in each hemisphere (Dice, 1945; Shrout and Fleiss, 1979).

ICC(3,k) was computed for intra-rater reliability (consistency) and ICC(2,k) was computed for inter-rater reliability (agreement). Dice was derived using the formula 2*(intersecting region)/(original segmentation + repeat segmentation); a Dice overlap metric of 0 represents no overlap, whereas a metric of 1 represents perfect overlap. Intra-rater and inter-rater reliability results (Table 2) were comparable to reliability values reported in the literature for manual segmentation of hippocampal subfields and MTL cortices (Wisse et al., 2012; Yushkevich et al., 2015b) and are consistent with our previous work (Olsen et al., 2013; Palombo et al., 2013).

**Table 2.**
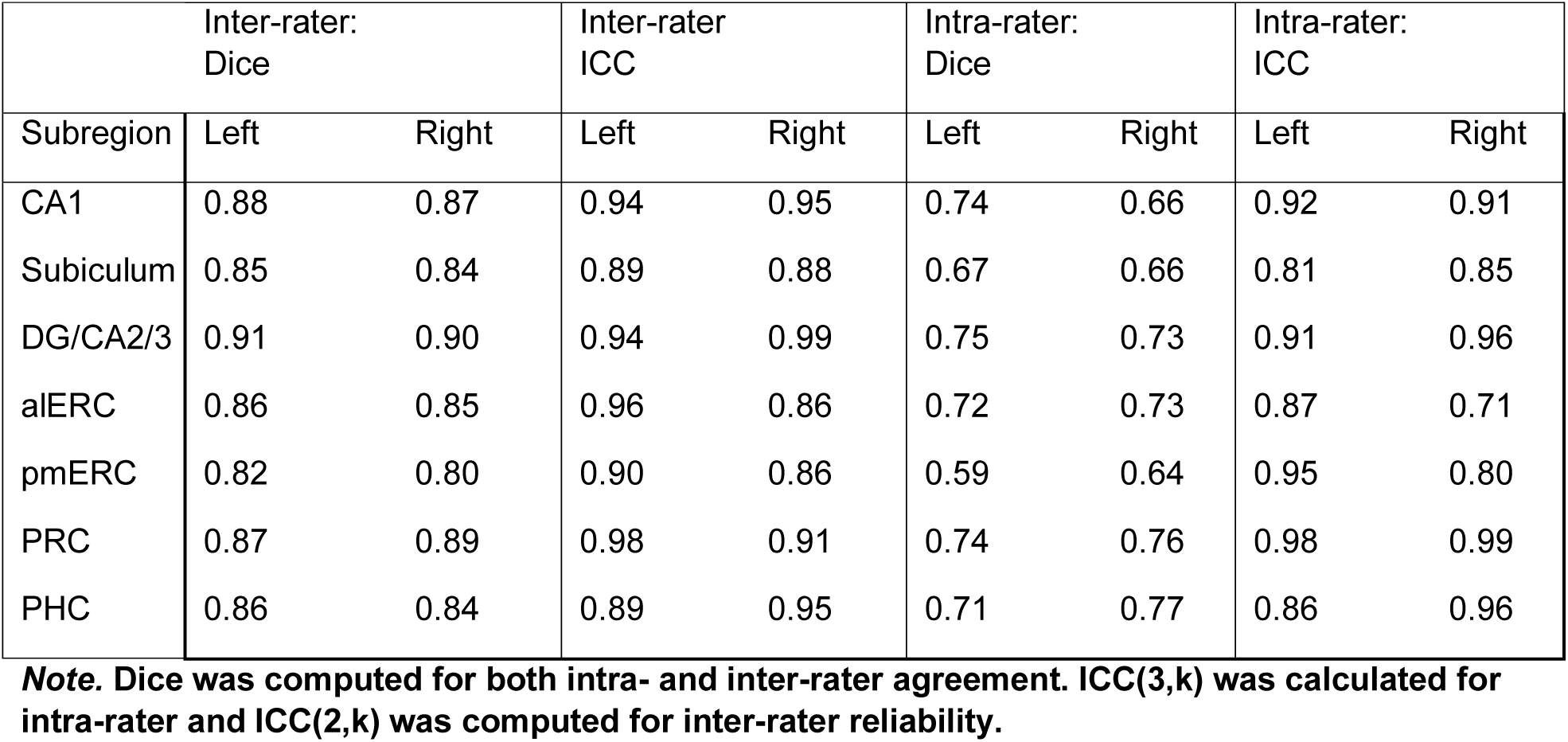
Reliability measurements.

#### Head size correction

All segmented volumes were corrected for head size to account for overall size differences across participants.(Arndt et al., 1991; Voevodskaya et al., 2014) By regressing the volume of each region with eTIV, a regression slope (β) was obtained for each region (representing the effect of eTIV change on that region’s volume). Then, the volume of each region was adjusted by that participant’s eTIV using the formula:

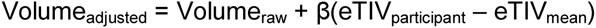

The head size correction was computed for each region separately for each hemisphere.

### 2.7 Statistics

Group differences were evaluated with t-tests and repeated measures ANOVAs in SPSS (version 22; IBM SPSS Statistics for Windows) Given the extensive literature reporting volume reductions in these regions as a function of AD severity (Adachi et al., 2003; Kerchner et al., 2012, 2010; La Joie et al., 2013; Mueller and Weiner, 2009; Mueller et al., 2010b, 2007; Pluta et al., 2012; Wisse et al., 2014; Yassa et al., 2010), and our previous work on a similar group of individuals who demonstrated neural and behavioural impairments (D’Angelo et al., 2016; Newsome et al., 2013, 2012; Yeung et al., 2013), we had strong a priori hypotheses that brain volumes would be smaller in the at-risk group; thus, one-tailed tests were used when comparing both global and MTL regions. The three hippocampal subfields and four MTL cortical subregions were entered into a single ANOVA model to test for main effects of group and group by region interactions; significant interactions were followed up with independent t-tests. The Holm-Bonferroni method was used to control familywise error rate when performing multiple comparisons. To characterize the nature of brain volume changes on cognitive performance, bivariate correlations were calculated between MoCA and the volume of brain regions that demonstrated significant group differences.

## 3 Results

### 3.1 Global Neuroimaging Measurements

Estimates of eTIV, cortical grey matter volume, cortical white matter volume, and CSF for each group were compared for each group. No significant group differences were observed for the cortical grey matter and CSF volume measures, and only a marginal difference was observed for cortical white matter (Supplementary Table 1; *p*s>.08). WMH volume for each group was also examined and a marginal group difference was observed (t(38)=1.48, *p*=.08). Visual examination of the FLAIR images ruled out the presence of previously undetected stroke.

### 3.2 Group Differences in MTL Subregion Volumes

A repeated-measures ANOVA was performed with brain region as a within-subjects factor, and group [at-risk, healthy] as a between-subject factor. Initial exploration of the data revealed no significant group X hemisphere interactions; consequently, the reported analyses were run on left and right hemispheres averaged. Mauchly’s test indicated that the assumption of sphericity had been violated for the effect of brain region (χ^2^(2)=279.52, *p*<.001). Therefore, degrees of freedom were corrected using Greenhouse-Geisser estimates of sphericity (ε=0.30).

There were significant main effects of brain region, F(6,228)=404.27, *p*<.001, η^2^=0.91, and group, F(1,38)=6.19, *p*<.001, η^2^=.14, and a significant brain region x group interaction, F(6,228)=2.57, *p*=.04, η^2^=.06.

The mean volumes (and SD, in mm^3^) of each of the three hippocampal subfields and four MTL cortex subregions, in the at-risk and healthy groups are listed in Supplementary Table 2; boxplots for each region are plotted in Figure 2. Follow-up independent samples t-tests showed that only the alERC region was significantly larger in the healthy versus the at-risk group (t(38)=3.37, *p*=.001), when accounting for multiple comparisons. The CA1 subfield (t(38)=2.40, *p*=.01), the perirhinal cortex (t(38)=2.04, *p*=.02) also showed group differences; however, these effects did not survive correction for multiple comparisons.

**Figure 2.**
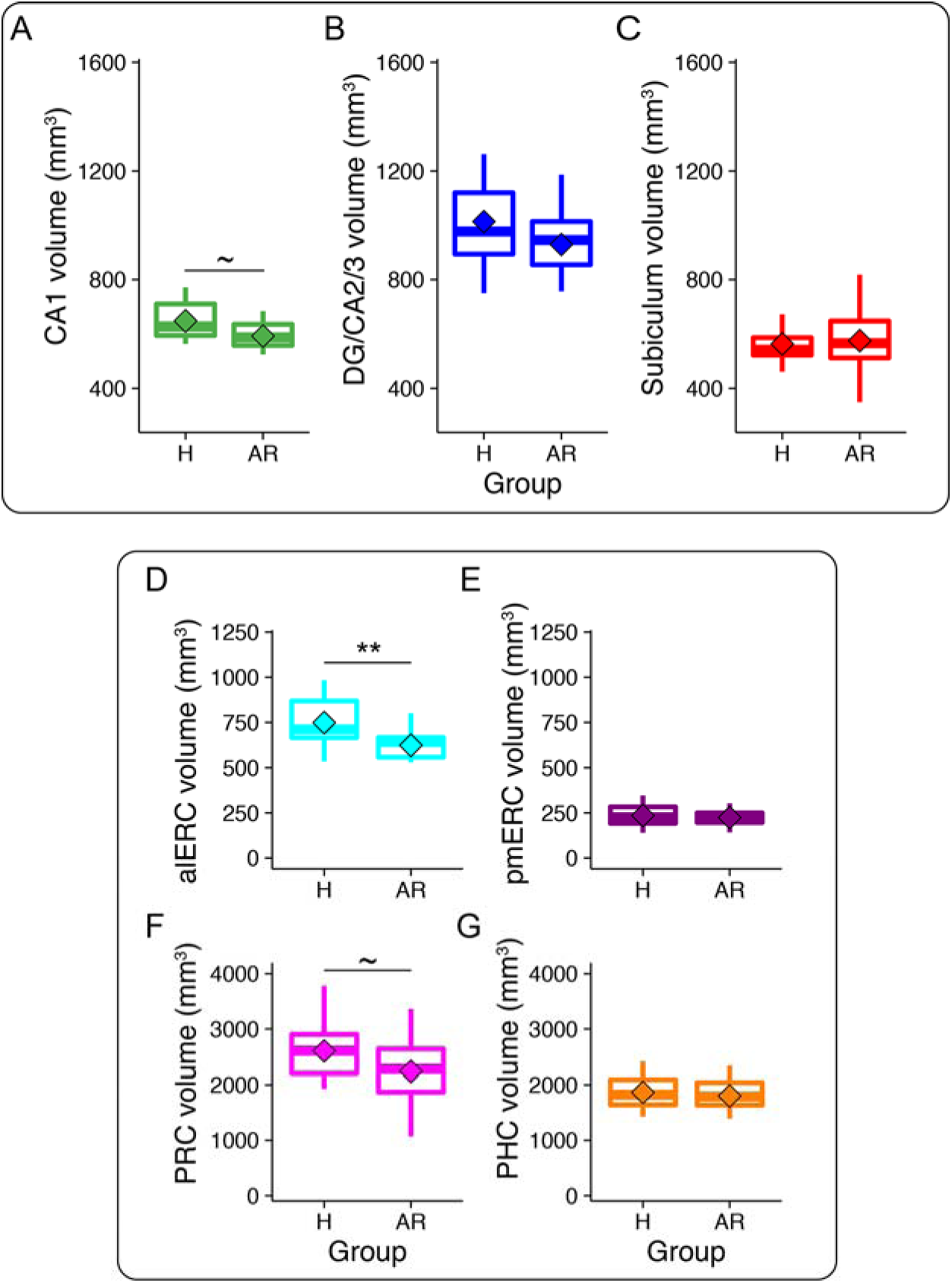
Hippocampal subfields and MTL cortical volumes. Box plots, plotted separately for healthy (H) and at-risk (AR) participants, for the hippocampal subfields (A-C) and MTL cortical subregions (D-G). ~Indicates a significant difference at p < .05 (does not survive multiple comparisons), ** p < .01.

### 3.3 Relationship Between Overall MoCA Performance and alERC Volumes Across Participants

In order to evaluate the nature of the relationship between structural atrophy in the alERC and severity of cognitive impairment, a bivariate correlation between MoCA score and alERC volume was conducted. This analysis showed a significant correlation between MoCA score and alERC volume (r=0.37, p=.009). Larger volumes in the alERC were associated with higher MoCA scores (Figure 3); this relationship remained significant even after accounting for any effect of age (r=0.34, p=.02).

**Figure 3.**
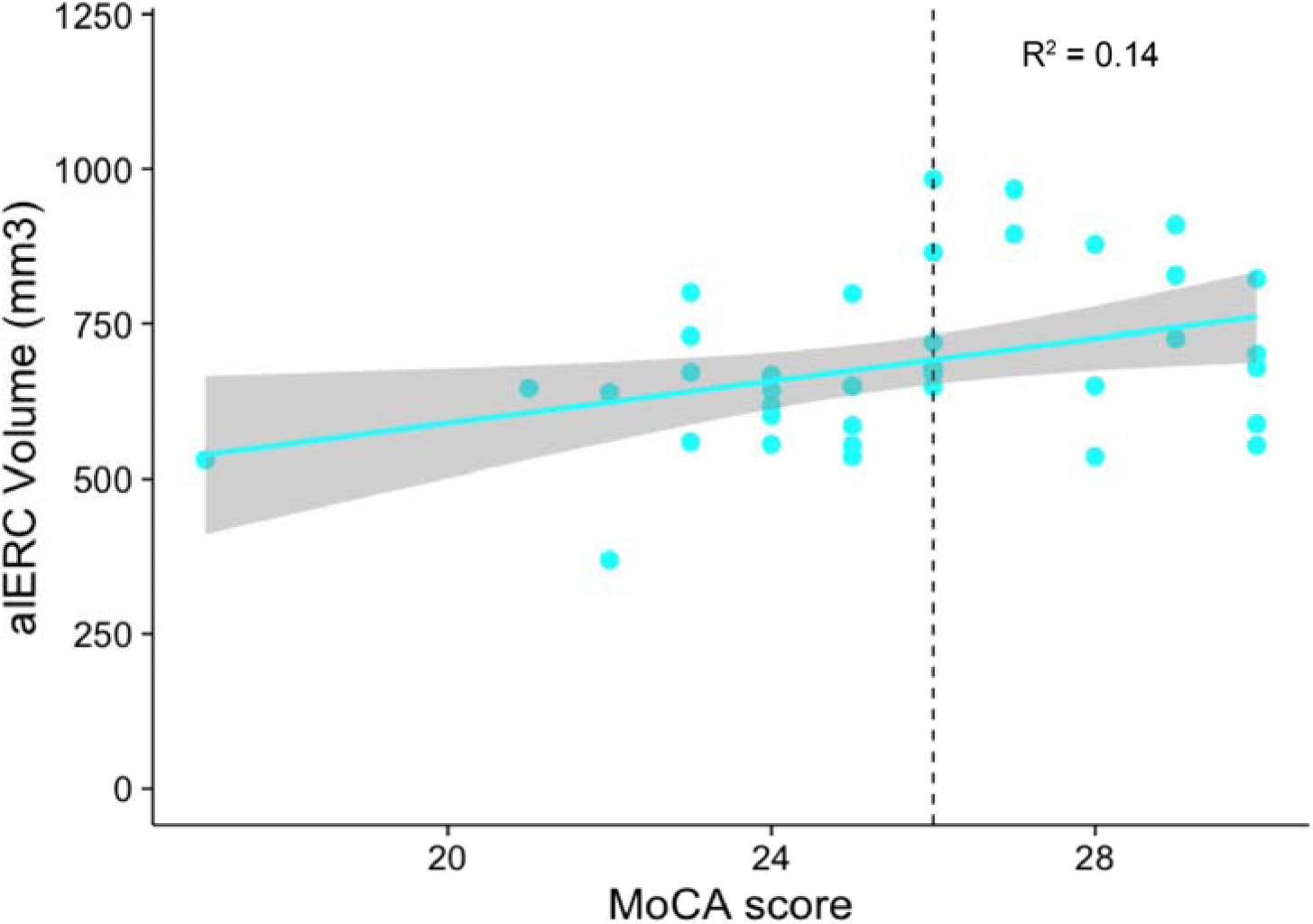
MoCA score correlates positively with alERC volume. Scatter plot depicting the nature of the relationship between alERC volumes and MoCA scores (each point reflects a single participant). Dashed line indicates the cut-off score that differentiated the healthy and at-risk groups. The shaded region indicates the 95% confidence region predicted by the linear model.

## 4. Discussion

We investigated MTL subregion volumes associated with preclinical cognitive decline in an ostensibly cognitively healthy older adult group. These individuals reported no subjective memory complaints but nonetheless scored below the recommended MoCA threshold, indicating that these individuals were at risk for developing dementia. Compared to the healthy group, we found a significant volume reduction for the at-risk group in the alERC, a region implicated in the early stages of AD. The significant positive relationship between alERC volume and MoCA score was also observed when MoCA was treated as a continuous, rather than a categorical variable, and when controlling for the effect of age. These results are important and novel for two reasons.

First, to our knowledge, this is the first study to employ segmentation of the alERC as a distinct region from the ERC as a whole, and thus we provide novel findings that link alERC volume to cognitive impairment. Combined with recent research suggesting that AD pathology originates from the alERC (Khan et al., 2014), and the extensive body of work which has shown overall volume reductions in the ERC as a function of MCI or AD status (Augustinack et al., 2012; Bobinski et al., 1999; Du et al., 2003, 2001; Fennema-Notestine et al., 2009; Fjell et al., 2014; Fujishima et al., 2014; Juottonen et al., 1998; Kerchner et al., 2013; Killiany et al., 2002; Mcdonald et al., 2009; Pennanen et al., 2004; Wisse et al., 2014), we propose that low alERC volume may be an early biomarker for AD risk. While the MoCA results and selective alERC volume reductions in our at-risk group suggest that these individuals may have asymptomatic, preclinical AD, this cannot be resolved with certainty without testing for the presence of AD biomarkers (i.e. amyloid-beta or tau level in the CSF, PET scan with amyloid-binding contrast) and/or a formal neuropsychological diagnosis with longitudinal follow-up. Further prospective/longitudinal studies are necessary to determine whether alERC volume is a sensitive and specific marker for AD in currently preclinical individuals.

Second, this is the first study to directly show that lower MoCA scores are related to volumes in specific MTL subregions that are affected in AD. While the MoCA has been shown to have a high specificity and selectivity for cognitive impairment, only two studies have previously examined the relationship between the MoCA and brain volume (Gupta et al., 2015; Paul et al., 2011) and neither of these studies looked specifically at MTL cortical subregions or hippocampal subfields. Our findings show that reduced alERC volumes, and to a lesser extent, reduced CA1 and PRC volumes, precede subjective memory complaints in community dwelling individuals, and further support the use of the MoCA as a predictive measure for AD (Julayanont et al., 2014; Nasreddine et al., 2005).

Recent work examining the relationship between global neuroimaging measurements and MoCA score reported significant associations between overall grey matter volume and CSF with cognitive performance (Gupta et al., 2015). In the current study, however, global neuroimaging measurements did not demonstrate significant group differences.

In conclusion, this is the first study to show reduced alERC volumes in ostensibly cognitively healthy, preclinical individuals who scored poorly on the MoCA, suggestive of AD-related cognitive decline, in the absence of any group differences in global brain volume or WMH. Importantly, the alERC is the brain region in which AD pathology is believed to originate (Yassa, 2014), and the reductions observed here may reflect early AD pathology. This research reveals mechanisms underlying pathological aging, and provides a potential neural target for early screening, evaluation of disease progression, and intervention efficacy.

## Acknowledgments

This work was supported by grants from the Canadian Institutes of Health Research (CIHR) to MDB (grant number MOP-115148) and JDR (grant number MOP-126003). MDB and JDR are supported by Canada Research Chairs. MDB is also supported by a Scholar Award from the James S McDonnell Foundation. L-KY is supported by a Natural Sciences and Engineering Research Council (NSERC) Canadian Graduate Scholarship.

